# The Role of Secretome from Mesenchymal Stromal Cells in Promoting Nerve Regeneration After Neurotmesis

**DOI:** 10.1101/2025.07.24.666642

**Authors:** Yaiza González Rodríguez, Alejandro Casado Santos, María Rodríguez-Díaz, Endika Nevado-Sánchez, Francisco Isidro Mesas, Irene Martín-Tamayo, Susana Martínez-Flórez, María Luisa González-Fernández, Jorge Labrador-Gómez, Vega Villar-Suárez

## Abstract

**Introduction:** Neurotmesis, the most severe form of peripheral nerve injury, remains a significant clinical challenge due to limited intrinsic regenerative capacity and suboptimal outcomes of current therapies. Mesenchymal stromal cells (MSCs) secretome has emerged as a promising cell-free alternative, providing neurotrophic and immunomodulatory factors to support nerve repair while avoiding the limitations of cell transplantation. This study aimed to evaluate the regenerative efficacy of primed adipose-derived MSC secretome in a rat model of sciatic nerve neurotmesis.

**Materials and Methods:** Human and rat adipose-derived MSCs were cultured and primed under hypoxic and inflammatory conditions. Secretomes were characterized by nanoparticle tracking analysis, proteomics, and total protein quantification. Neurotmesis was induced in Wistar rats, followed by repair with biomaterial alone or combined with human or rat secretome. Functional recovery was assessed by neurophysiological measurements (CMAP, NAP) at 6 months. Molecular and protein analyses included qPCR for myelination genes and ELISA for NGF. Morphological regeneration was evaluated by histology, immunofluorescence, and transmission electron microscopy (TEM).

**Results:** Secretome priming enhanced the secretion of neurotrophic factors (GDNF, VEGFA, FGF2) and immunomodulatory proteins (IL6, CCL2), as confirmed by transcriptomic and proteomic analyses. *In vivo*, secretome-treated groups showed significantly improved neurophysiological recovery, with restoration of CMAP/NAP amplitudes and increased NGF levels. qPCR revealed upregulation of myelination-associated genes (*MPZ, Krox-20, c-Jun*) in treated nerves. Histological and TEM analyses demonstrated robust axonal regeneration, thicker myelin sheaths, and the presence of Remak bundles.

**Conclusions:** Primed MSC secretome markedly enhances structural and functional recovery after sciatic nerve neurotmesis, supporting its potential as a safe, effective, and scalable cell-free therapy for peripheral nerve repair. These findings provide a strong rationale for further translational studies and clinical development.

## 1. Introduction

Neurotmesis represents a severe form of peripheral nerve injury (PNI) that involves significant challenges for recovery due to the limited intrinsic repair mechanisms of the nervous system ^1^. These injuries often result in severe functional deficits, impacting on the quality of life of affected individuals. The process of nerve regeneration involves a series of cellular and molecular events, including inflammation, axonal sprouting, and remyelination, which are crucial for restoring nerve function ^2^. Schwann cells (SCs) play a pivotal role in the peripheral nervous system (PNS), facilitating nerve regeneration by producing neurotrophic factors and supporting axonal growth ^3^. However, in cases of severe nerve damage, such as neurotmesis, the natural repair process is often insufficient, necessitating therapeutic interventions. Current treatments for PNI include surgical repair, physical therapy, and pharmacological interventions, but these methods have limitations in terms of efficacy and speed of recovery ^4^.

Recent advancements in tissue engineering and regenerative medicine have highlighted the potential of mesenchymal stem/stromal cells (MSCs) in promoting nerve regeneration ^5–8^. MSCs are an ideal cell source for tissue regeneration due to their outstanding properties. They are multipotent stromal cells capable of differentiating into various cell types, including adipocytes, osteoblasts, chondrocytes, myocytes, β-pancreatic islet cells, and potentially neuronal cells ^9^. However, recent studies have shown that implanted cells have a limited survival time, and their clinical implementation faces challenges such as immunological incompatibility, tumour formation, and potential infection transmission ^10^. New findings have highlighted the diverse range of bioactive factors produced by MSCs, which may play a crucial role in regulating various physiological processes.

Therefore, the secretome from MSCs has attracted significant attention for its potential use in tissue repair and regeneration^11–13^. The secretome is a collection of bioactive molecules and factors secreted by cells into the extracellular space ^14,15^. It includes proteins, growth factors, cytokines, chemokines, and extracellular vesicles that transport lipids, proteins, and various types of RNA and DNA. These paracrine factors can prevent apoptosis, promote cell proliferation, and stimulate neovascularization, providing essential support for damaged tissues. The secretome-based therapy offers several advantages over traditional cell therapy ^16–18^. Being cell-free, it reduces the risks of immune rejection, tumorigenicity, and pathogen transmission, while simplifying storage and administration. Additionally, the secretome can be produced in large quantities and stored for extended periods without the need for toxic cryoprotectants, making it more scalable and accessible for clinical applications. This approach allows for the delivery of therapeutic factors without the challenges associated with whole cell therapies, providing a promising tool for tissue repair and regeneration ^19–22^.

An important consideration in the therapeutic application of MSCs is the concept of “priming.” In their native state, MSCs exhibit significant heterogeneity, which can lead to variability in their therapeutic effects. Priming refers to the process of preconditioning MSCs to enhance their inherent properties and tailor their therapeutic potential to specific clinical needs ^23^. This involves exposing MSCs to certain environmental cues or biochemical signals that optimize their secretory profile and improve their ability to respond to tissue damage. By mitigating the variability in MSC responses, priming aligns with the principles of personalized medicine, allowing for customized therapeutic outcomes that meet the unique requirements of individual patients and medical conditions. This approach has the potential to reveal new possibilities in regenerative medicine by enabling the precise modulation of MSCs functions to address specific clinical challenges. Previous studies in our lab have focused on establishing a priming protocol for adipose tissue-derived MSCs, aiming to identify candidates with optimal anti-inflammatory and regenerative properties ^24^.

The primary goal of this work is to provide scientific evidence on the efficacy of stromal cell secretome in promoting peripheral nerve regeneration. The results of this study could have significant clinical implications, contributing to the development of safer and more effective nerve regeneration therapies that avoid the challenges associated with whole cell therapies. Specifically, we propose using the primed secretome from adipose tissue-derived mesenchymal stromal cells (ASCs) to assess its potential in regenerating sciatic nerve neurotmesis in a rat model. This approach aims to serve as a complementary therapeutic strategy to existing treatments, enhancing the outcomes of conventional methods.

## 2. Materials and methods

### 2.1. Cell culture and Secretome Collection

The MCB-HTC-22 LP01 ASCs line (Histocell^®^) derived from human adipose tissue was cultured in MEM Alpha medium (MEM α Gibco^®^), supplemented with 3% platelet lysate (PL, Elarem^®^), 1% Penicillin-Streptomycin (Pen Stre, Gibco^®^), 0.1% Amphotericin (HyClone^®^), 0.1% Gentamicin (Gibco^®^), 0.02% Vancomycin (Thermo Scientific^®^), and 0.75 units/mL of heparin (Sigma-Aldrich^®^) to create a complete growth medium. All experiments were performed using cells at passages 2 to 4. Cells were maintained in a humidified environment with 5% CO_2_ at 37°C. Secretome from rat ASCs was also used as control. Rat ASCs were obtained as described by González-Cubero et al. ^8^.

For secretome production, cells were primed under hypoxic and inflammatory conditions, after which conditioned media were collected, filtered (0.22 µm) (Merck Millipore^®^), and stored at −80°C.

### 2.2. Transcriptomic Analysis

Primed cells were collected and subjected to NGS analysis to examine genes that were upregulated or downregulated in ASCs following priming conditions. Prior to sequencing, RNA extraction and quality was assessed with the Quant-iT RiboGreen RNA Assay Kit (Invitrogen^®^) with a VICTOR Nivo Multimode Microplate Reader (PerkinElmer^®^), employing fluorescence-based quantification via Quant-iT™ RiboGreen RNA Reagent^®^. RNA quality was evaluated by assessing its integrity with the Agilent 2100 Bioanalyzer Instrument (Agilent^®^).

Libraries were prepared and sequenced, with pseudo-alignment and quantification performed using Salmon (GRCh38 reference genome ^25^). Differential expression analysis was carried out in DESeq2 package ^26^, with significance set at padj< 0.05 and log□ fold change of ±1. Enrichment of WikiPathways and Gene Ontology terms was evaluated using Gene Set Enrichment Analysis (GSEA) algorithm ^27^.

### 2.3. Secretome characterization

Total protein content was quantified using the Micro BCA Protein Assay Kit (Thermo Scientific^®^). Nanoparticle tracking analysis (NTA) was performed with a NanoSight LM10^®^ system to determine particle size and concentration.

For proteomic analysis, secretome proteins were acetone-precipitated, digested via the FASP method ^28^, and analyzed by LC-MS/MS on a timsTOF HT mass spectrometer coupled to an Evosep ONE liquid chromatograph. Protein identification and quantification were conducted using Fragpipe software against the Mus musculus Uniprot/Swissprot database, with a 1% FDR cutoff. Only proteins present in ≥70% of samples per group were included for further analysis. Missing values were imputed (QRILC method, Perseus v.2.1.2.0 ^29^.), statistical significance was assessed by t-test (p<0.05) and the data are represented as log2 fold change.

### 2.4. *In vivo* model of sciatic nerve neurotmesis

#### 2.4.1. Animals

The experimental protocols in this study conform to the guidelines established by the Directive 2010/63/EU and adhere to Spanish regulations (RD 53/2013) for laboratory animal use. The experiments were performed in line with the standards set by the relevant Spanish regulations and the ethical guidelines of the OEBA (Animal Welfare Body) and the OH (Authorized Body) of HUBU and authorized by the Junta de Castilla y León. In this study, forty female Wistar rats (*Rattus norvegicus*), aged 5 to 10 months and weighing 300 to 400 grams were used. These animals were maintained in conventional housing conditions: a temperature of 22 ± 2°C, relative humidity of 55 ± 10%, and a 12-hour light-dark cycle. A standard diet was provided with *ad libitum* access to food and water. Measures were taken to minimize animal suffering and limit the number of animals used.

#### 2.4.2. Surgical procedure

Forty female Wistar rats (5–10 months, 300–400 g) were housed under standard conditions. Under a combination of ketamine (Ketolar^®^, 50 mg/ml) at 60 mg/kg and xylazine (Xilagesic^®^, 20 mg/ml) at 7.5 mg/kg administered intraperitoneally, the sciatic nerve was exposed and completely transected (neurotmesis) 10 mm distal to its origin, followed by epiperineural suture with 10-0 nylon. Experimental groups included: (1) control (contralateral limb), (2) neurotmesis + biomaterial, (3) neurotmesis + biomaterial + human secretome, and (4) neurotmesis + biomaterial + rat secretome. Secretome was delivered both via a pericardium membrane (Jason^®^) wrapped around the lesion and by intraneural injection. The biomaterial group received only the membrane (Figure 1A). Skin was closed with 4-0 nylon. Animals were sacrificed at 6 months post-surgery for tissue analysis using qPCR, histology, TEM, confocal microscopy and ELISA assays.

**Figure 1:**
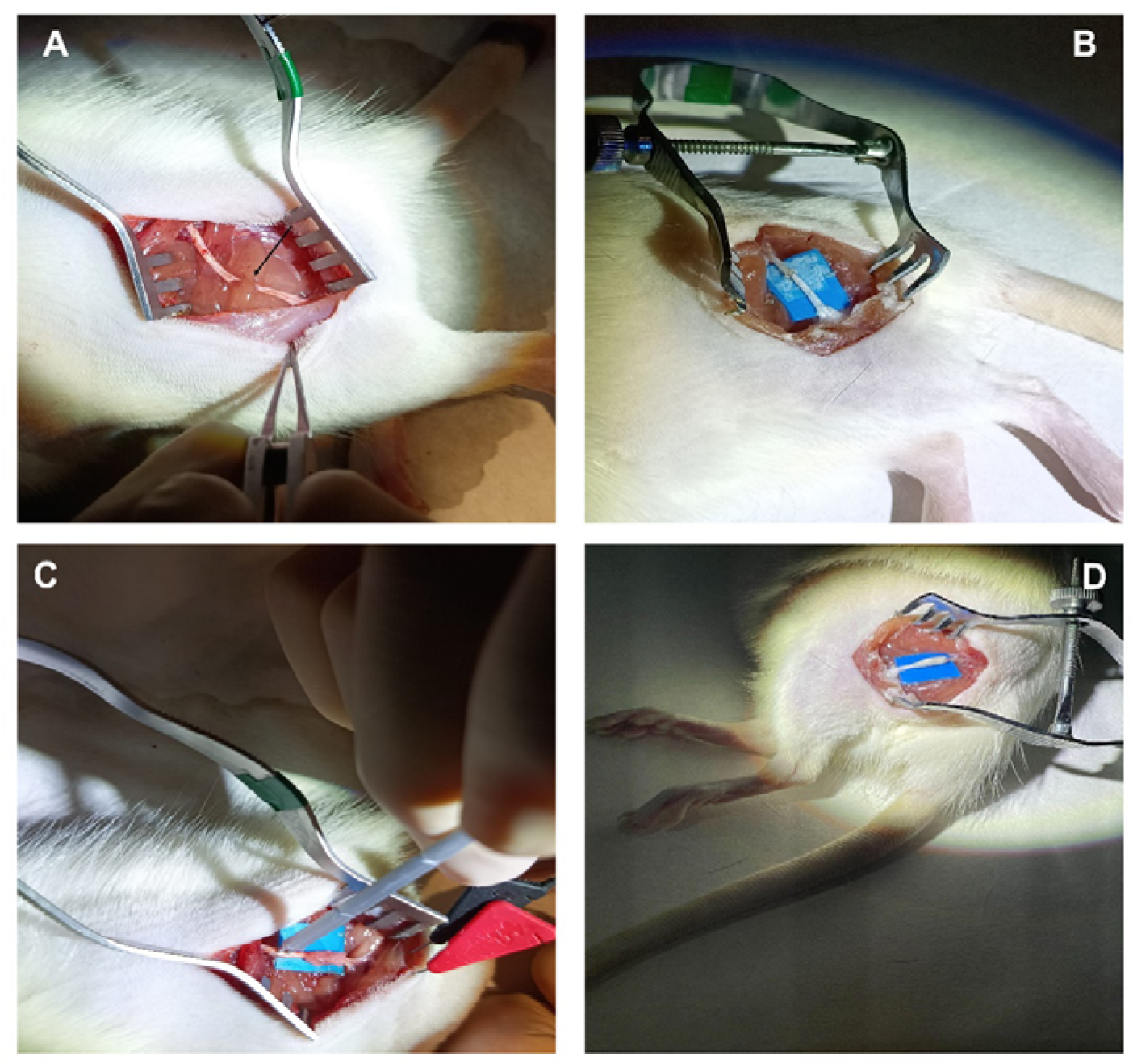
Surgical procedure. (A) Neurotmesis of sciatic nerve (arrow). (B) Sciatic nerve sutured. (C) Biomaterial placed below the sutured sciatic nerve. (D) Biomaterial embedded in secretome sleeve around the sciatic nerve.

#### 2.4.3. Neurophysiological analysis

This assessment was performed after surgical exposure of the target nerves by generating a nerve action potential (NAP) and a compound muscle action potential (CMAP), both before and after nerve transection (neurotmesis) and suturing, intraoperatively. During the follow-up period, each individual will undergo evaluations at 12 weeks post-intervention and prior to animal sacrifice (after 6 months), to identify signs of active reinnervation or functional recovery. These evaluations will utilize quantitative electromyography (EMG) and electroneurography (ENG), using a Natus Nicolet Synergy EDX EMG System^®^. Statistical analyses were performed using R Studio and IBM SPSS Statistics (v20), with a significance level set at 0.05 (95% confidence interval). Qualitative variables were described as absolute frequencies and percentages, with group comparisons assessed using Fisher’s exact test for 2×2 tables and the Freeman-Halton extension for variables with more than two categories. For ordinal or non-parametric variables, the Mann-Whitney U test was used for multiple comparisons. Normality of continuous variables was evaluated with the Kolmogorov-Smirnov and Shapiro-Wilk tests. For the analysis of neurophysiological potentials (NAP and CMAP), paired Student’s t-tests were applied to compare pre- and post-intervention values within each group, assessing both latency and amplitude; results were reported as mean differences, standard deviation (SD), standard error, 95% confidence intervals, and corresponding p-values.

#### 2.4.4. qPCR

Total RNA was isolated from nerve tissue using GeneMATRIX Universal RNA Purification Kit and quantified with a Qubit 4 Fluorimeter. cDNA synthesis was performed with a high-capacity kit (Applied Biosystems^®^). qPCR was run using Power SYBR Green PCR Master Mix on a StepOne real-time PCR system, with GAPDH and ACT-β as reference genes. The 2^-ΔΔCt^ method was used for quantification. Specific primers (Table 1) were designed using the Primer-BLAST tool and synthesized by Sigma-Aldrich^®.^ Three biological replicates per condition were analyzed, with results expressed as mean ± SD. ANOVA and Tukey’s post-hoc test were used for statistical analysis (p<0.05).

**Table 1:**
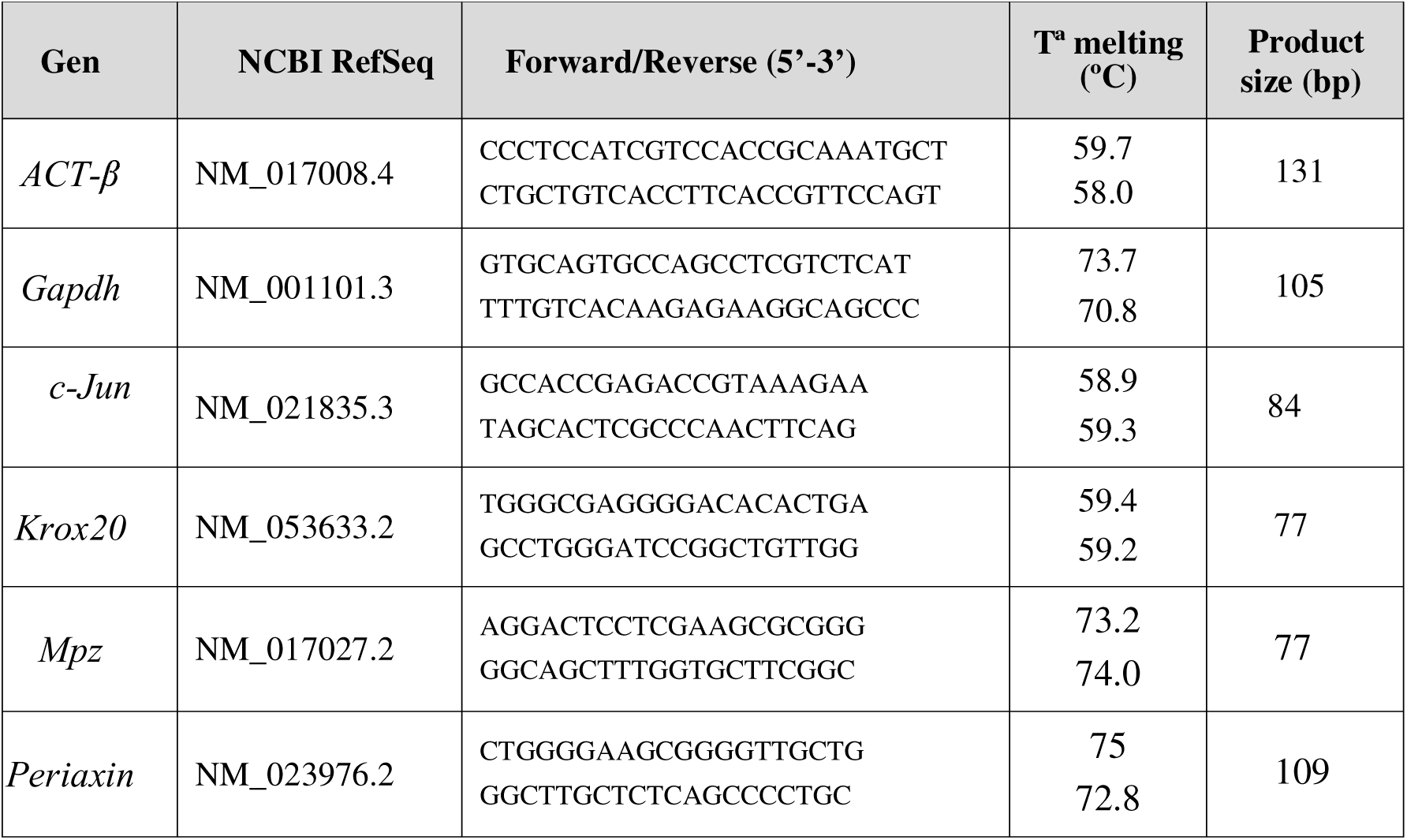
Primer sequences and conditions used for qPCR.

#### 2.4.5. ELISAs

Frozen tissue samples (−80°C) were pulverized using a liquid nitrogen-prechilled metal pulverizer (Spectrum™). ELISA assay for Nerve Growth Factor (NGF) (Cusabio^®^) was subsequently performed according to the manufacturer’s instructions. Statistical analysis was conducted as per the methods outlined for qPCR and therefore omitted to avoid redundancy.

#### 2.4.6. Histology

Nerve samples were fixed in 4% paraformaldehyde, dehydrated, and embedded in paraffin. Sections (2 μm) were stained with hematoxylin and eosin for general morphology. Semi-thin sections (0.5 μm) were stained with toluidine blue for axon quantification. Four random fields per sample were analyzed using a Nikon E600 microscope with NIS Elements software^®^. Statistical analysis also followed the approach described for qPCR.

#### 2.4.7. Confocal microscopy

For immunofluorescence, sections were blocked and incubated with anti-MPZ primary antibody (1:100, Invitrogen^®^) overnight at 4°C, followed by Alexa Fluor 488-conjugated secondary antibody (1:100, Invitrogen^®^) and DAPI nuclear counterstain. Slides were mounted with Fluoromount. Imaging was performed using a Zeiss^®^ confocal microscope and analyzed using the NIS Elements microscope imaging software (Nikon^®^). Statistical analysis also followed the approach described for qPCR.

#### 2.4.8. Transmission Electron Microscopy (TEM)

Nerve ultrastructure was assessed using TEM. Nerves were fixed in 2.5% glutaraldehyde, post-fixed in 2% osmium tetroxide, dehydrated, and embedded in epoxy resin. Ultra-thin sections (60 nm) were cut and stained with uranyl acetate and lead citrate, then examined with a JEM-1010 JEOL^®^ TEM. Ten random fields per group were analyzed for axon and myelin measurements (area, diameter, circularity, G-ratio, myelin width, axon/sheath ratio) using NIS Elements microscope imaging software (Nikon^®^). Measurements from the inner axons and myelin sheaths were acquired independently, including area (µm2), circularity and equivalent diameter (µm). Derived measurements (G-ratio, myelin sheath width (µm), and Axon/Sheath ratio) were calculated from primary measurements.

Morphometric analysis results are presented as mean ± SD. For primary measurements, data normality was assessed, and either one-way ANOVA or Kruskal–Wallis test was applied based on the outcome. For derived measurements, either standard one-way ANOVA or Brown–Forsythe and Welch ANOVA was used, depending on the homogeneity of variances. Post hoc analyses were conducted for multiple group comparisons where appropriate. Results with p<0.05 were considered statistically significant. GraphPad Software^®^ was used for statistical analysis.

## 3. Results

### 3.1. Gene Ontology Enrichment Analysis Reveals Pro-Regenerative Transcriptional Profile

Gene ontology (GO) enrichment analysis revealed significant upregulation of biological processes associated with nerve regeneration and repair (Figure 2A). Key enriched categories included neural development, motor neuron axon guidance (normalized enrichment score ∼1.8), and positive regulation of neurogenesis. Among upregulated genes (Figure 2B), critical neurotrophic factors showed substantial expression changes: glial cell-derived neurotrophic factor (*GDNF*, log2 fold change ∼3.5), vascular endothelial growth factor A (*VEGFA* ∼3.0), fibroblast growth factor 2 (*FGF2* ∼2.8), and insulin-like growth factor 2 (*IGF2* ∼6.0), all promoting neuronal survival and axonal outgrowth. The regeneration-associated activating transcription factor 3 (*ATF3*) was also significantly upregulated (∼3.0 fold change).

**Figure 2:**
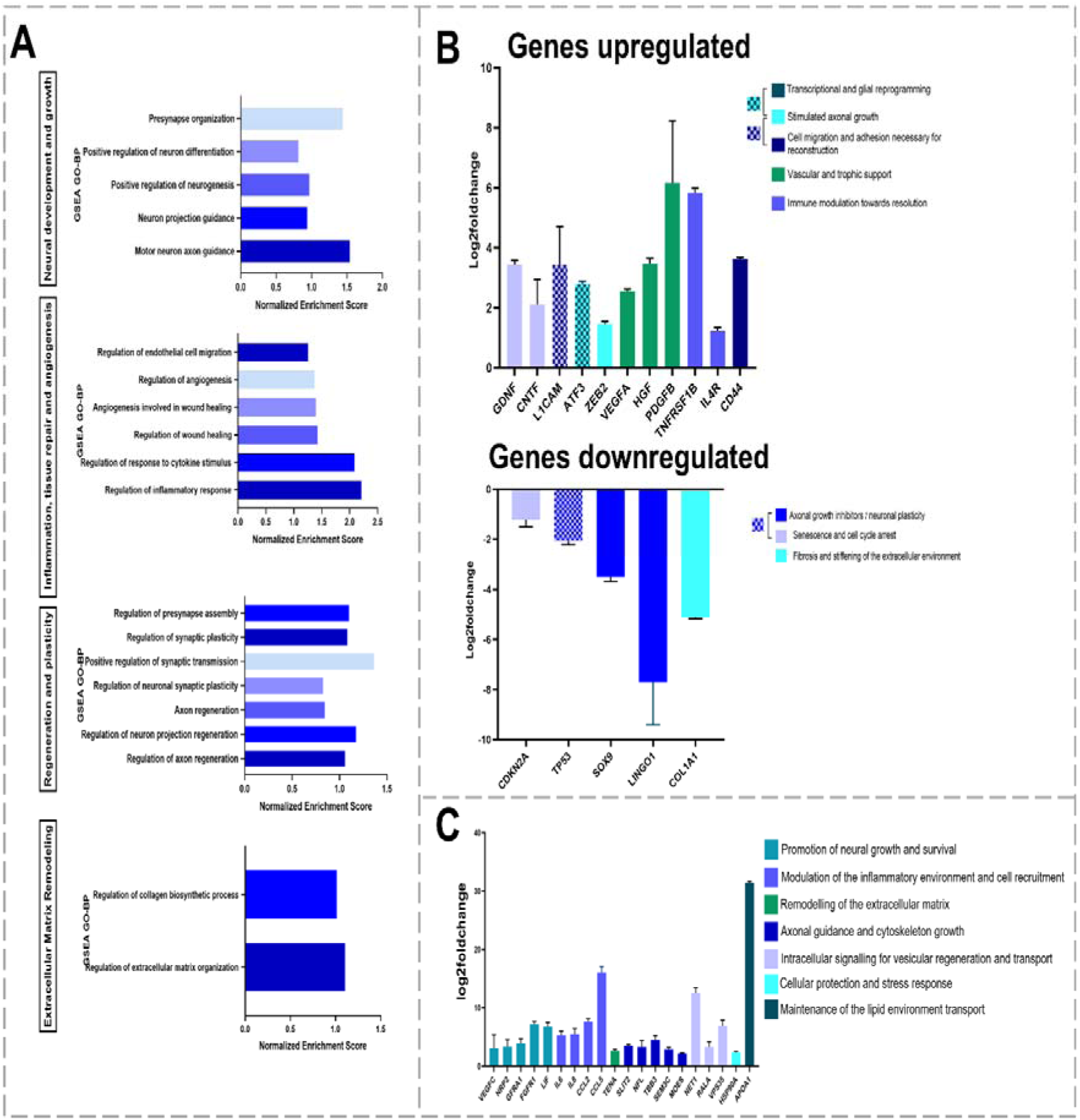
Transcriptomic changes in licensed mesenchymal stem cells reveal a pro-regenerative profile relevant to sciatic nerve regeneration. (A) Gene set enrichment analysis (GSEA) of differentially expressed genes. (B) Bar plots showing representative upregulated (top) and downregulated (bottom) genes related to nerve regeneration. (C) Quantitative proteomic analysis of selected secretome proteins involved in nerve regeneration. Data are presented as mean ± SEM.Data are presented as log2 fold change ± SEM.

Immune modulation genes including tumor necrosis factor receptor superfamily member 1B (*TNFRSF1B* ∼1.5 fold change) and *IL1R* (interleukin 1 receptor ∼3.8 fold change) were coordinately upregulated (Figure 2B), alongside extracellular matrix remodeling processes that facilitate axonal regeneration. Conversely, several axonal growth inhibitors were significantly downregulated, most notably leucine-rich repeat and immunoglobulin-like domain-containing no-go receptor-interacting protein 1 (*LINGO1* ∼-8.0 fold change), tumor protein p53 (*TP53* ∼-2.0 fold change), and cyclin-dependent kinase inhibitor 2A (*CDKN2A* ∼-1.5), removing major barriers to regeneration and potentially facilitating cell cycle re-entry.

This comprehensive transcriptional reprogramming demonstrates a coordinated shift toward a pro-regenerative phenotype, characterized by enhanced neurotrophic support, reduced growth inhibition, and active tissue remodeling—all essential components for successful peripheral nerve regeneration.

### 3.2. Secretome characterization

Table 2 presents secretome characterization comparing standard normoxic (2D normoxia) and priming conditions (2D hypoxia inflammation). NTA analysis revealed no significant differences in particle concentration (∼1.6×10^9^ vs. ∼1.5×10^9^ particles/mL) or average particle size (∼160 vs. 158 nm) between conditions. However, hypoxic-inflammatory priming significantly increased total protein concentration (p<0.01) from ∼1.4 to ∼1.55 μg/mL, indicating enhanced protein content despite consistent particle numbers. This suggests that while the particle number remains consistent, the protein content within these particles or in the soluble fraction is enhanced under inflammatory and hypoxic stimulation.

**Table 2:** Characterization of secretome under normoxic and hypoxia/inflammation conditions.

| Sample | N° Particle/mL |  |  | Size (nm) |  |  | Protein concentration (Mg/mL) |  |  |
| --- | --- | --- | --- | --- | --- | --- | --- | --- | --- |
|  | Mean | SD | p value | Mean | SD | p value | Mean | SD | p value |
| **NORMOXIA** | 1,60E+09 | 1,73E+08 | 0,519 | 165,57 | 7,20 | 0,384 | 1,41 | 0,01 | 0,023 |
| **HYPOXIA**<br>**INFLAMMATION** | 1,50E+09 | 1,73E+08 |  | 161,43 | 1,37 |  | 1,54 | 0,03 |  |

Proteomic analysis (Figure 2C) revealed distinct enrichment of nerve regeneration-associated proteins. Key neurotrophic factors (VEGFC, NRP2, GFRA1, FGFR1, LIF) supported neuronal survival and axonal growth, while inflammatory modulators (IL6, IL8, CCL2, CCL5) facilitated reparative cell recruitment. Additional proteins supported axonal guidance and cytoskeleton dynamics (TENA, SLIT2, NEFL, TBB3, NET1), intracellular signaling for vesicular transport (RALA, VPS35, HSP90AA), and lipid environment maintenance (APOA1) for membrane repair and remyelination. These findings demonstrate that primed secretome contains multiple orchestrated factors facilitating a comprehensive peripheral nerve repair.

### 3.3. *In vivo* assessment of nerve regeneration 6 months after surgery

The primary aim of our research was to evaluate the potential of human secretome derived from human ASCs in nerve regeneration. To provide a comparative context and assess the efficacy of the human secretome, we also used rat secretome as a control in our experimental model. This approach allowed us to assess whether there were significant differences in the outcomes between human and rat secretome, while accounting for interspecies differences.

#### 3.3.1. ASC secretome restores ANP and CMAP to near baseline levels

Neurophysiological evaluation (Figure 3) demonstrated that secretome treatment, particularly with human-derived secretome, resulted in markedly superior functional recovery compared to control and biomaterial-only groups. In the control group 75% of animals exhibited absent or reduced NAP (Figure 3A), and 87.5% showed degraded, polyphasic CMAP (Figure 3B), reflecting poor reinnervation and severe functional deficit. The biomaterial-only group demonstrated some improvement, with all animals presenting PAN and PAMC, though 37.5% remained degraded and EMG parameters were only modestly improved. In contrast, the human secretome group showed robust recovery (Figure 3C), with 100% of animals displaying strong, morphologically valid PAN and PAMC (78.9% high-quality signals, (Figures 3A and 3B), and EMG recordings indicating advanced stages of active reinnervation. The rat secretome group also exhibited substantial improvement, though with slightly fewer motor unit potentials than human secretome group. Quantitatively (Figure 3D), this human secretome group demonstrated significantly higher EMG amplitude compared to neurotmesis and biomaterial (p<0.001), and rat secretome group (p<0.05). Duration and polyphasia were also significantly greater in the human secretome group versus control (p<0.001) and biomaterial (p<0.05) (Figure 3D). The sum of these parameters, reflecting advanced reinnervation, was significantly higher in the human secretome group compared to all other groups (p<0.001 vs. biomaterial; p<0.01 vs. rat secretome). The contraction pattern was also most favorable in this group, with significant differences compared to neurotmesis (p<0.01), biomaterial (p<0.05), and rat secretome groups (p<0.01) (Figure 3D). The percentage recovery of NAP amplitude was highest in the human secretome group, with significant differences compared to untreated (p<0.01) and biomaterial groups (p<0.05) (Figure 3C). Finally, the distribution of global classification scores by group is illustrated in the bubble plot (Figure 3E), further supporting the superior neurophysiological recovery achieved with human secretome treatment. Collectively, these results demonstrate that ASC secretome, especially of human origin, markedly enhances neurophysiological recovery and active reinnervation after sciatic nerve injury.

**Figure 3:**
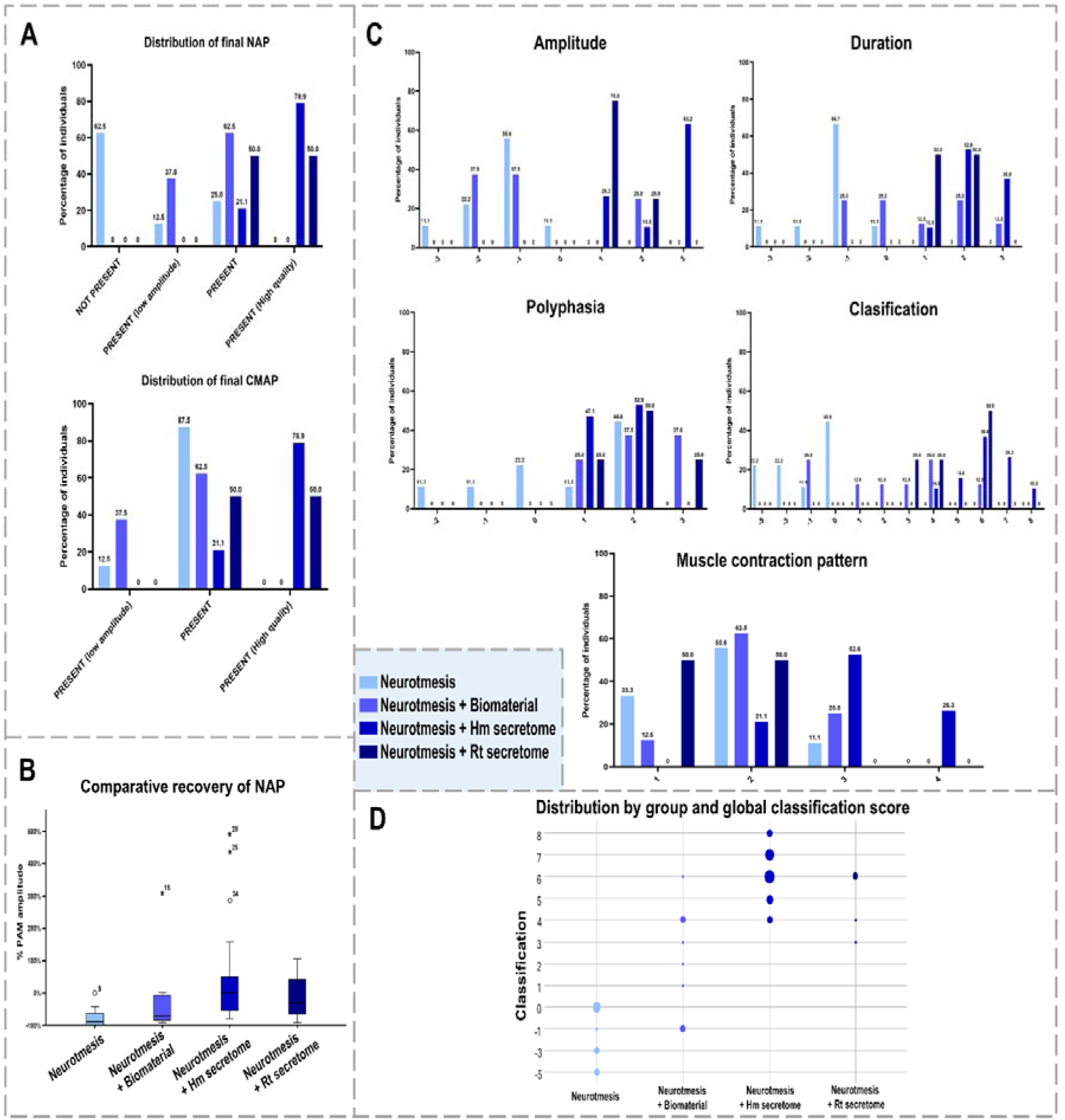
Neurophysiological outcomes by experimental group. (A) Distribution of animals by final NAP status. (B) Distribution by final CMAP status. (C) Comparative boxplot of NAP amplitude recovery between groups. (D) Percentage distribution of EMG parameters: amplitude (Amp), duration (Dur), polyphasia (Ppp), global classification (Clasif), and contraction pattern (Pattern) by group. (E) Bubble plot showing the distribution of animals by group and global classification score.

#### 3.3.2. ASC secretome upregulates myelination-inducing genes and nerve-specific proteins in axonal repair

Figure 4A illustrates relative mRNA expression of four critical myelination-associated genes (*MPZ*, *KROX-20*, *c-JUN*, and *Periaxin*) in rat sciatic nerve following different experimental conditions. Our qPCR analysis revealed striking expression patterns across experimental groups. In the pro-myelination markers *MPZ*, *KROX-20* and *c-JUN*, biomaterial application with suture repair demonstrated remarkable upregulation (∼ 3.5-4 fold change increase compared to control), suggesting potent regenerative activation. Interestingly, human secretome treatment maintained moderately raised expression levels of these genes, while rat secretome application resulted in significant suppression. Conversely, Periaxin expression exhibited an inverse pattern, with substantial downregulation across all intervention groups compared to control, being most pronounced in the neurotmesis group. The neurotmesis with suture-only group consistently showed inhibited expression of MPZ and c-JUN compared to control, indicating impaired natural regenerative capacity.

**Figure 4:**
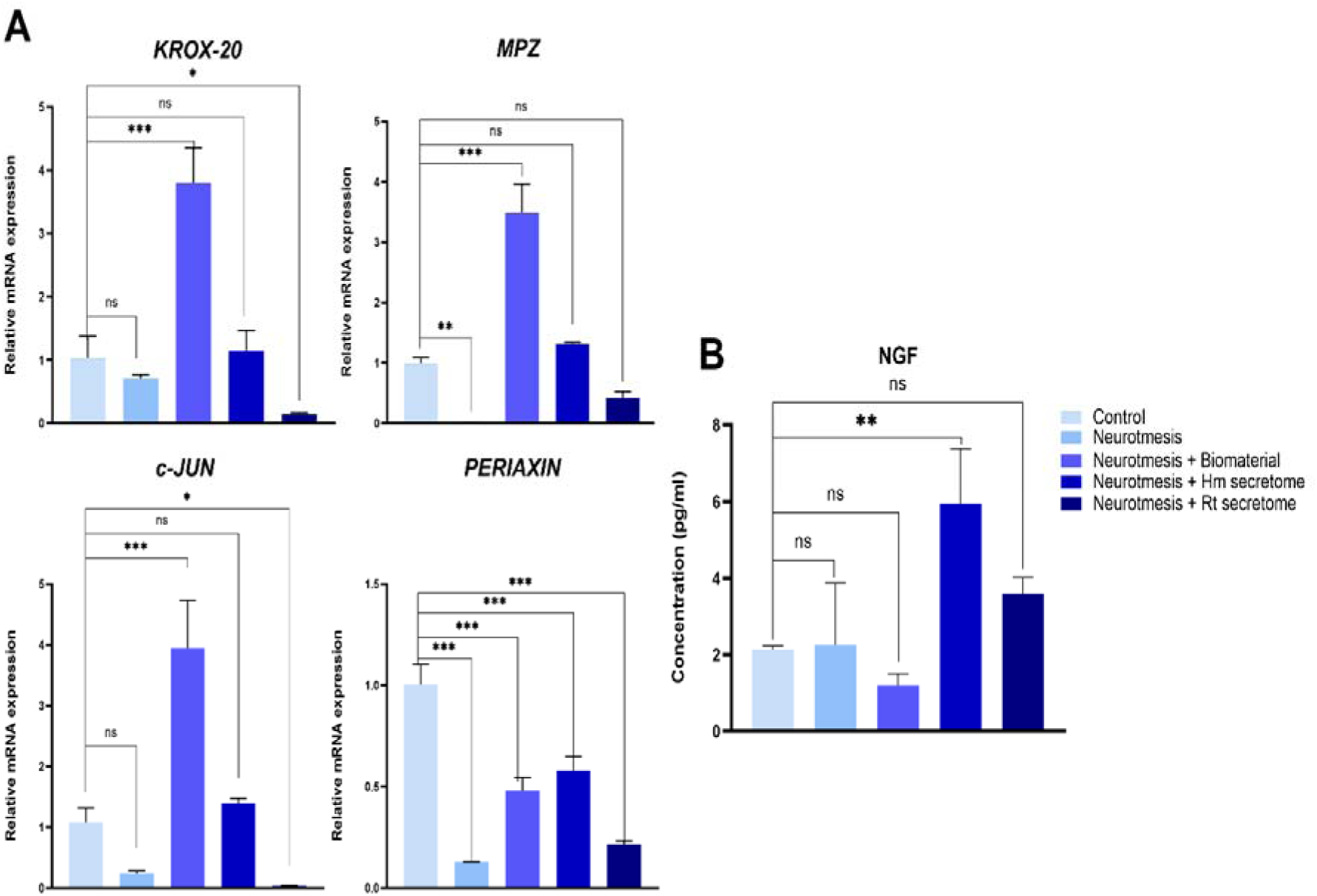
Gene expression analysis and NGF protein quantification. (A) Quantitative PCR analysis of key genes involved in nerve regeneration and myelination. (B) ELISA quantification of NGF protein levels. Data are presented as mean ± SEM. Statistical significance: *p<0.05, **p<0.01, ***p<0.001; ns, not significant.

The ELISA results for NGF concentration across experimental groups are shown in Figure 4B. No significant differences were observed between the control, neurotmesis, and neurotmesis plus biomaterial groups. However, the group treated with human secretome exhibited a significant increase in NGF levels compared to all other groups (p<0.01). The group treated with rat secretome also showed elevated NGF concentrations, although this increase was not statistically significant relative to the other conditions. These findings indicate that the application of the human secretome markedly enhances NGF expression at the injury site, suggesting a potent neurotrophic effect.

#### 3.3.3. ASC secretome promotes remyelination in treated nerves

##### Longitudinal sections highlight segmental differences in nerve architecture

Longitudinal sections of the sciatic nerve stained with hematoxylin and eosin (Figure 5) reveal distinct histological differences among the experimental groups. The control group displays the typical architecture of healthy nerve tissue, with highly organized, wavy fascicles, compact parallel fibers, and low cellularity. In contrast, the neurotmesis group exhibits partial disruption of fascicular organization, irregular fiber alignment, and moderate cellular infiltration, reflecting tissue injury and early regeneration. The biomaterial group shows increased cellularity with numerous elongated nuclei and evidence of neovascularization (white arrows), as well as the presence of macrophage-like cells (asterisks). Notably, both the human and rat secretome-treated groups demonstrate further enhancement of regenerative features, including a higher density of blood vessels (white arrows) and abundant macrophage-like cells (asterisks), supporting robust angiogenesis and debris clearance.

**Figure 5:**
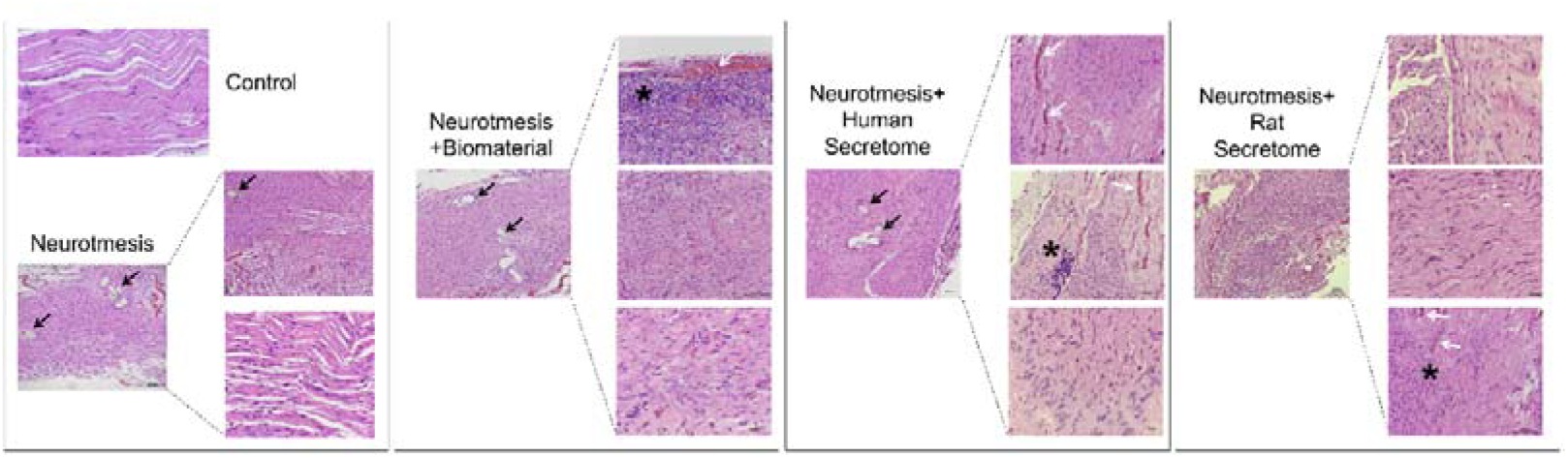
Representative longitudinal sections of sciatic nerve stained with hematoxylin and eosin. From left to right: healthy control (up), neurotmesis with suture (down), neurotmesis with suture and biomaterial, neurotmesis with suture and human secretome, and neurotmesis with suture and rat secretome. Suture zone (black arrows); blood vessels (white arrows), macrophages (white arrows). Scale bar: 20 μm and 100 μm.

Immunofluorescence analysis of longitudinal sciatic nerve sections stained for MPZ (Figure 6) revealed clear architectural differences between groups, with the control nerves displaying highly organized, densely packed myelinated fibers, while neurotmesis and biomaterial groups showed disrupted fascicular structure and reduced axonal organization, especially in distal segments. Both human and rat secretome treatments improved axonal density and alignment, partially restoring the fascicular architecture towards that of healthy nerves. However, despite these structural differences, no significant changes in MPZ fluorescence intensity were observed among any of the experimental groups, including neurotmesis, compared to controls.

**Figure 6:**
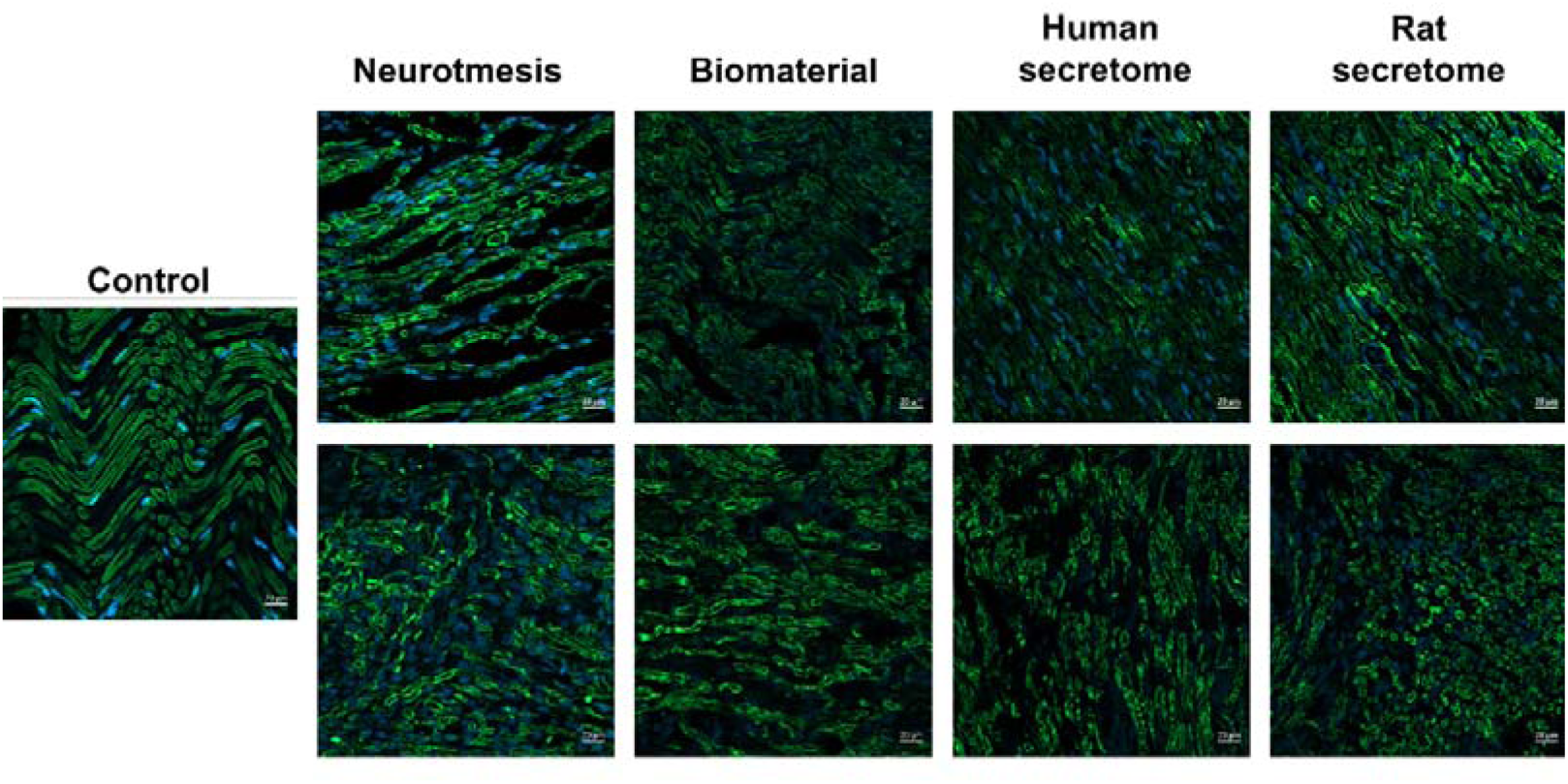
Representative longitudinal sections of sciatic nerve stained by immunofluorescence. (green: axonal/myelin marker; blue: nuclei) from each experimental group. Top row: proximal segments to the lesion site; bottom row: distal segments. Scale bar: 20 μm.

##### Toluidine blue staining highlights structural restoration and regeneration across treatments

Semithin sections of sciatic nerve stained with toluidine blue (Figure 7) reveal clear differences in axonal morphology and nerve architecture among groups. The control group (Figure 7 A.1-A.2) displays compact, highly organized fascicular structures with densely packed, uniformly sized myelinated axons. In contrast, the neurotmesis group (Figure 7 B.1-B.2) shows marked disorganization, loss of fascicular architecture, reduced axon density, and large endoneurial spaces, indicative of severe degeneration. The biomaterial group (Figure 7 C.1-C.2) presents partial preservation of axonal profiles and some improvement in myelinated fiber density compared to neurotmesis alone, though organization remains irregular and axon diameters are heterogeneous. Both the human secretome (Figure 7 D.1-D.2) and rat secretome (Figure 7 E.1-E.2) groups demonstrate substantial enhancement in nerve regeneration, with increased density of myelinated axons, more uniform diameters, thicker myelin sheaths, and partial restoration of fascicular organization. Notably, large new blood vessels (Bv) are observed in several treatment groups, supporting active regeneration. Quantitative analysis (Figure 7 F) confirms that all treatment groups (biomaterial, human secretome, rat secretome) have significantly higher axon counts than both control and neurotmesis groups (p<0.01, p<0.001), in direct correlation with the histological improvements seen in panels Figure 7 A–E.

**Figure 7:**
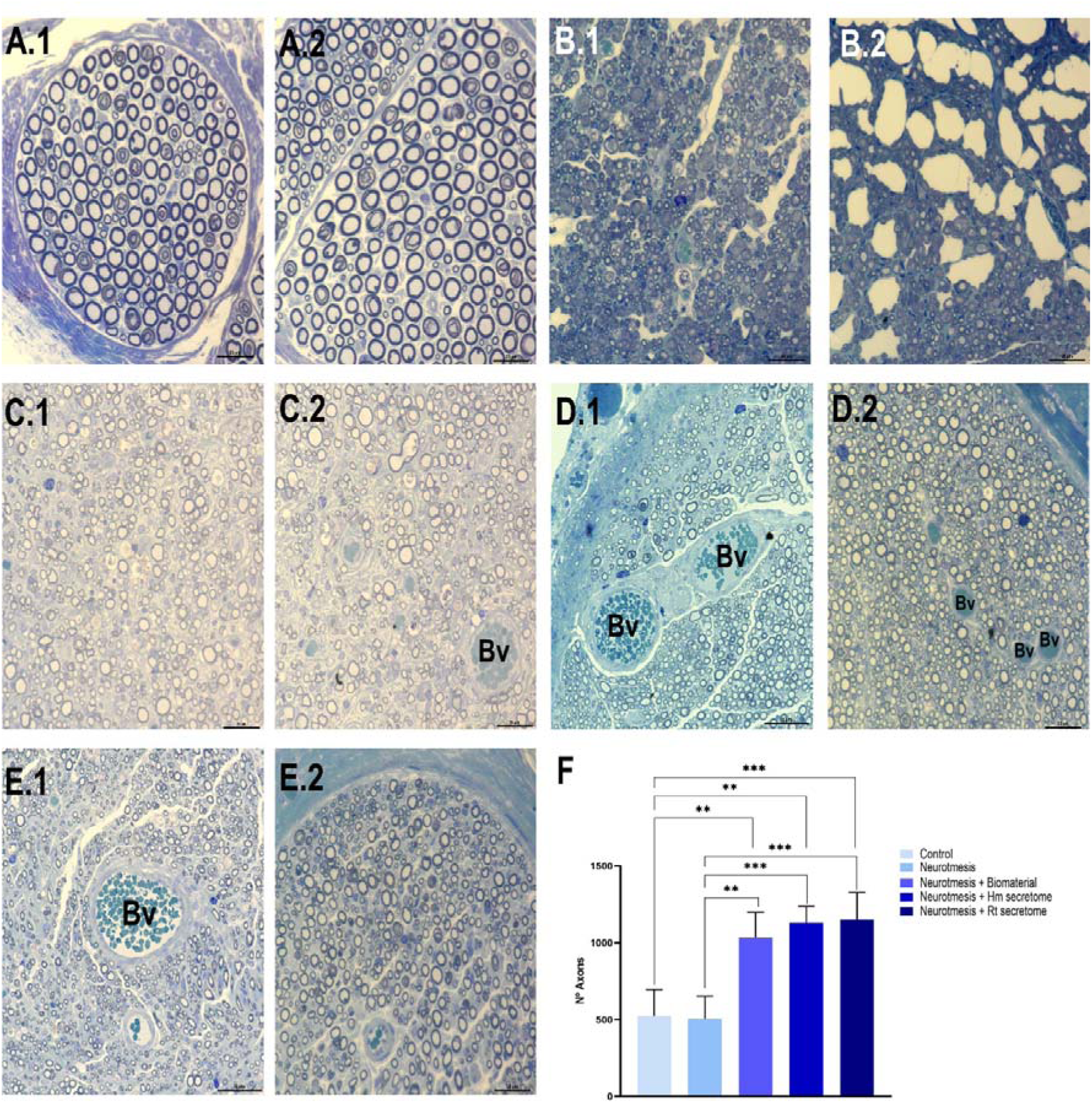
Representative semithin longitudinal sections of sciatic nerve stained with toluidine blue. (A.1, A.2) Healthy control. (B.1, B.2) Neurotmesis with suture. (C.1, C.2) Neurotmesis with suture and biomaterial. (D.1, D.2) Neurotmesis with suture and human secretome. (E.1, E.2) Neurotmesis with suture and rat secretome. Bv: Blood vessels. Scale bar: 20 μm. (F) Quantification of number of axons. Statistical significance: **p<0.01, ***p<0.001.

##### Transmission Electron Microscopy Reveals Superior Nerve Regeneration in Secretome-Treated Groups

The TEM images (Figure 8 A–E.3) and corresponding quantitative analysis (Figure 9) reveal significant ultrastructural differences in axonal regeneration and nerve architecture among the experimental groups. The control group (Figure 8 A) exhibited typical healthy peripheral nerve ultrastructure, with large, uniformly myelinated axons, thick myelin sheaths, minimal endoneurial space, and high circularity. There are few unmyelinated axons (asterisks), and Schwann cells appear normal in size and morphology (SC). In contrast, the neurotmesis group (Figure 8 B) showed severe architectural disruption, marked by a reduction in myelinated axons and the presence of characteristic onion bulb (OB) formations, indicative of chronic nerve injury and repeated demyelination/remyelination cycles. These features are quantitatively supported (Figure 9) by a significant decrease in axon area not correlated with a proportional reduction in myelin sheath area, due to the frequent OB structures. Circularity, along with G-ratio, also appear significantly lower, indicating a pathological thickening of myelin sheath and structural disorganization (p<0.001 vs. control). The biomaterial group (Figure 8 C) showed partial preservation of myelinated fibers compared to neurotmesis, with intermediate improvements in axon area and circularity, though they remained below control values. Endoneurial space remains expanded, and some large SCs are visible, indicating ongoing regeneration but incomplete recovery. Statistically, this group demonstrates intermediate values for axon area and circularity, significantly improved compared to neurotmesis alone (p<0.001), yet still lower than control. Both secretome-treated groups show notable regenerative activity. The human secretome group (Figure 8 D1-D3) revealed active regeneration with pro-myelinating Schwann cells (SCm), functional blood vessels (BV), and Remak bundles (Rb), showing improved organization, more uniform myelin thickness, as well as significant increases in axon and sheath areas, along with improved circularity, with G-ratio values approaching control. Most remarkably, the rat secretome-treated group (Figure 8 E1-E3) achieved near-complete regeneration, displaying ultrastructural features almost indistinguishable from healthy controls. This group exhibited well-organized, large-caliber axons, robust myelin sheaths, appropriately ensheathed unmyelinated fibers within well-organized Remak bundles, extensive pro-myelinating Schwann cell activity, and optimal vascular supply. Quantitative morphometric analysis (Figure 9) strongly supported these observations, confirming significantly higher axonal, myelin sheath, and total fiber areas, along with the restoration of normal circularity values and myelin sheath width, thus demonstrating that this treatment not only promotes axonal regeneration but also facilitates the complete restoration of peripheral nerve microarchitecture.

**Figure 8:**
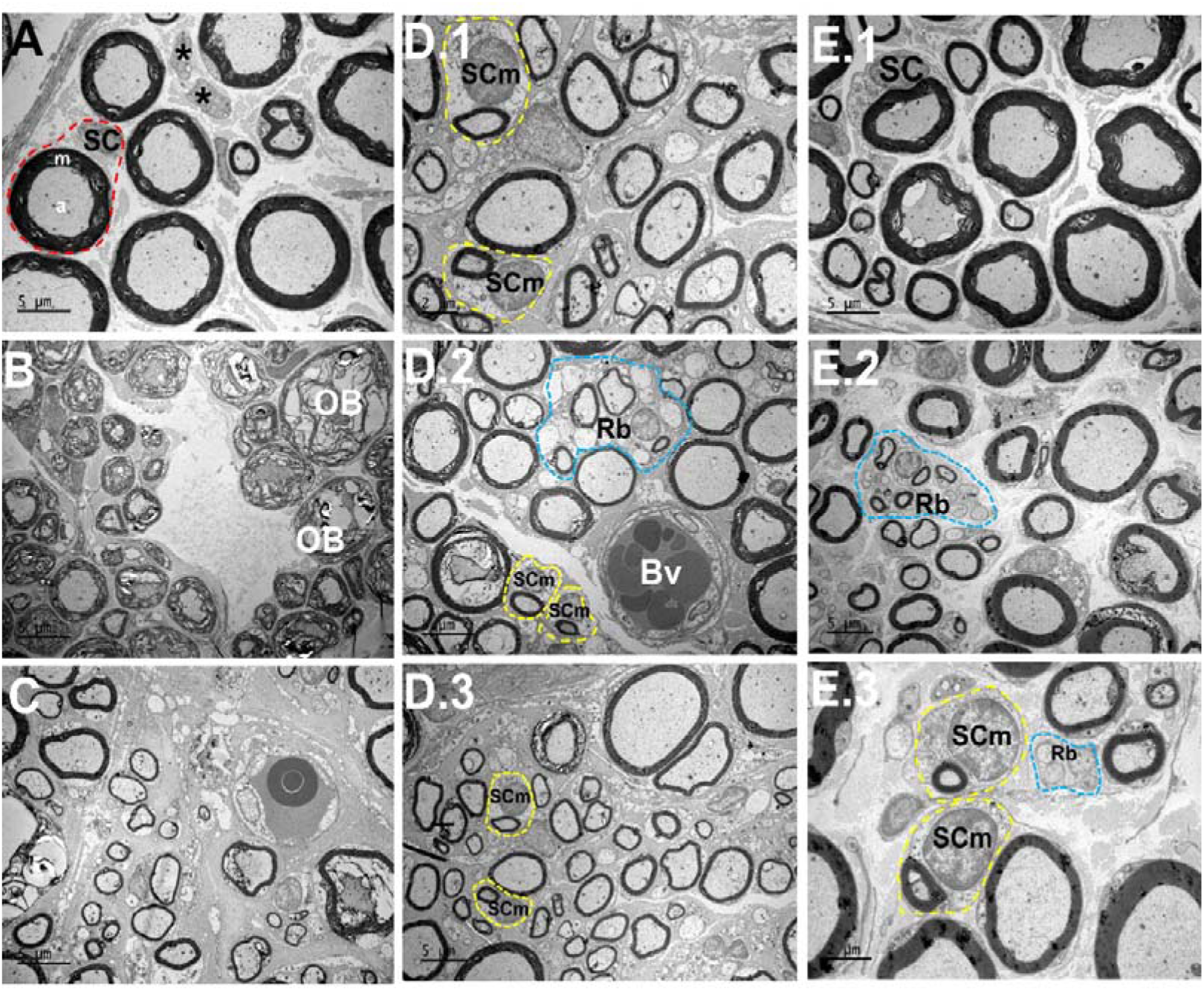
TEM images of sciatic nerve cross-sections from each experimental group. (A) Control nerve showing normal myelinated axons with compact myelin sheaths and organized fascicular structure. Schwann cells (SC), myelin sheath (m) and axon (a). (B) Neurotmesis group demonstrating severe nerve degeneration with disrupted myelin architecture, formation of onion bulbs (OB). (C) Neurotmesis treated with biomaterial showing partial structural recovery with some preserved myelinated fibers and activated Schwann cells. (D1-D3) Neurotmesis treated with human secretome displaying improved axonal organization, pro-myelinating Schwann cells (SCm), and presence of blood vessels (BV) indicating active vascularization. (E1-E3) Neurotmesis treated with rat secretome showing the most complete regeneration with well-organized myelinated axons, compact myelin sheaths, and Remak bundles (Rb). Scale bars = 5 μm. Scale bars: 5 μm (A, B, C, D.2, D.3, E.1, E.2); 2 μm (D.1, E.3).

**Figure 9:**
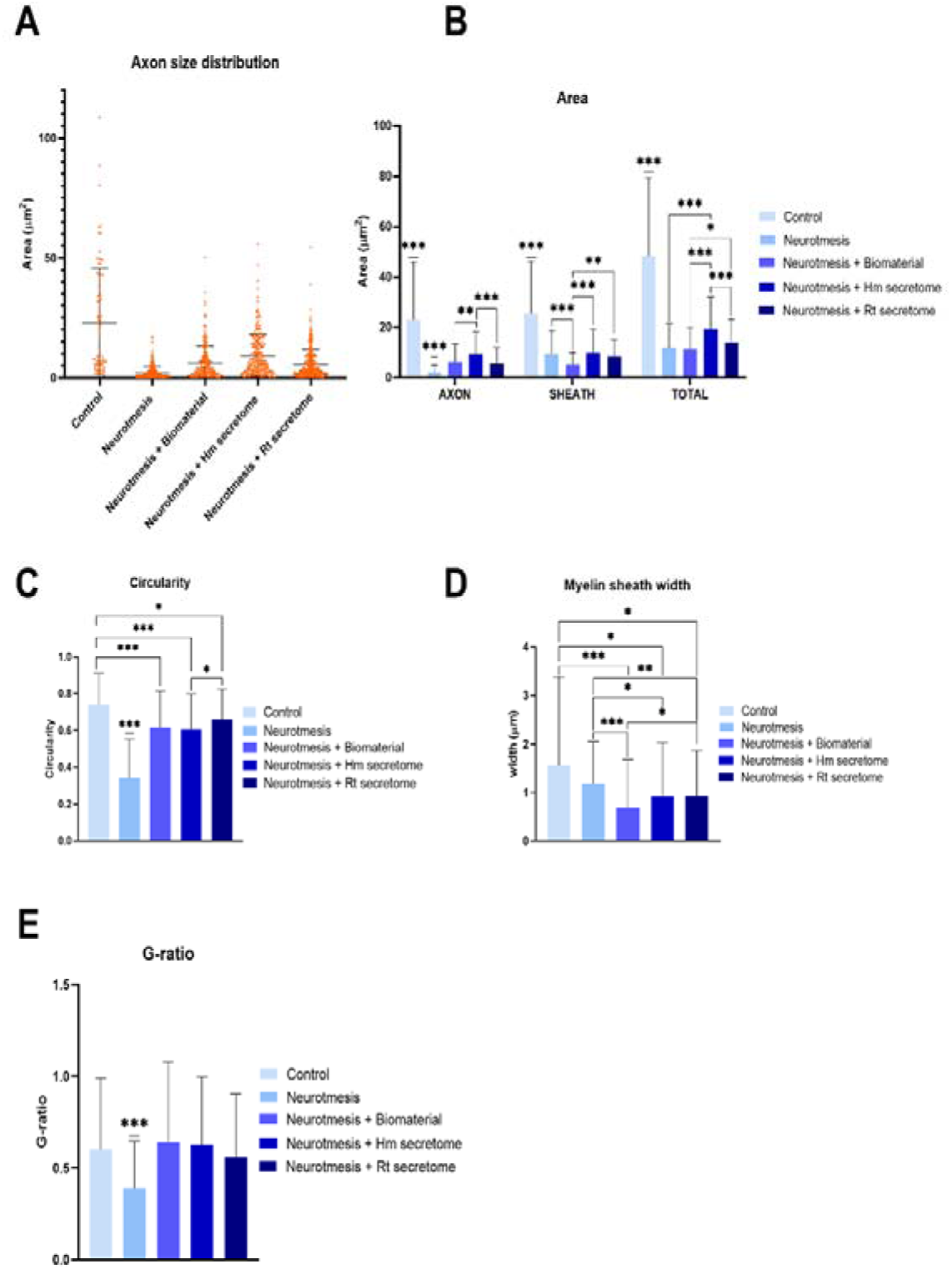
Quantitative morphometric analysis of nerve fiber regeneration parameters. (A) Axon size distribution across all experimental groups showing the range of axonal diameters. (B) Quantitative analysis of axonal area, myelin sheath area, and total fiber area demonstrating significant improvements in secretome-treated groups compared to neurotmesis alone (***p<0.001, **p<0.01, *p<0.05). (C) Circularity measurements indicating restored fiber geometry in treated groups, with significant differences between control and all injury groups (***p<0.001). (D) Myelin sheath width analysis showing progressive restoration in treatment groups, with significant differences between groups (*p<0.05, **p<0.01, ***p<0.001). (E) G-ratio analysis demonstrating optimal myelination patterns, with neurotmesis showing significantly altered G-ratio compared to control (***p<0.001). Data are presented as mean ± SEM. Statistical significance: *p<0.05, **p<0.01, ***p<0.001.

## 4. Discussion

This study aimed to advance the understanding and application of cell free therapy as a biological bridge for critical nerve lesions, specifically by researching the augmenting effect of ASC secretome on nerve regeneration and functional restoration. Our results indicate that optimized priming approaches (integrating hypoxia and inflammatory cues) substantially potentiate the therapeutic efficacy of ASC secretome by modulating their composition and functional properties.

The integration of transcriptomic and proteomic data provides compelling evidence that the ASC secretome promotes a comprehensive pro-regenerative response at the molecular, protein, and functional levels, thereby supporting robust peripheral nerve regeneration *in vivo*. Gene ontology enrichment analysis identified significant upregulation of biological processes directly associated with nerve regeneration, including neural development, axon guidance, and neurogenesis. Key neurotrophic factors such as *GDNF, VEGFA, FGF2*, and *IGF2* were markedly upregulated, supporting both neuronal survival and axonal outgrowth. These findings align with recent work by Kou et al. ^30^, who demonstrated that MSC-derived extracellular vesicles enhance axonal regeneration through similar neurotrophic factor delivery. Additionally, genes involved in immune modulation (e.g., *TNFRSF1B, IL1R*) and extracellular matrix remodeling were significantly activated, suggesting a coordinated molecular response that facilitates a regenerative microenvironment. Importantly, several inhibitors of axonal growth, such as *LINGO1* and *TP53*, were downregulated, further promoting a permissive state for nerve repair. This dual mechanism—enhancing growth signals while suppressing inhibitory pathways— endorses the therapeutic strategy proposed by Jessen and Mirsky ^31^, who emphasized the need to reprogram SCs by modulating both pro-regenerative and anti-regenerative signals. Our results agree with studies reporting limited efficacy of non-primed MSCs secretomes in nerve repair ^24,32^, highlighting the critical role of preconditioning strategies in optimizing secretome potency. Together, these findings primed ASC secretome as a multifaceted therapeutic agent capable of addressing the complex molecular challenges of peripheral nerve regeneration.

Proteomic profiling of the ASC secretome under both normoxic and hypoxic/inflammatory conditions demonstrated enrichment of proteins crucial for peripheral nerve regeneration. The secretome contained high levels of neurotrophic and growth-promoting factors (e.g., VEGFC, NRP2, GFRA1, FGFR1, LIF) that are essential for neuronal survival and axonal growth. This is consistent with previous reports that highlight the importance of such trophic factors in promoting axonal regeneration and functional recovery after nerve injury ^19^. Elevated concentrations of cytokines and chemokines (IL6, IL8, CCL2, CCL5) were observed, reflecting the secretome’s capacity to modulate inflammation and recruit reparative cells, in agreement with findings by Vizoso et al. ^16^. Proteins related to axonal guidance (TENA, SLIT2), cytoskeleton dynamics (TBB3, NEFL), and intracellular signaling (RALA, VPS35, HSP90AA) were also detected. The presence of APOA1 suggests an important role in maintaining the lipid environment for membrane repair and remyelination, a finding that aligns with studies emphasizing lipid metabolism as a determinant of efficient nerve regeneration ^33^. Altogether, these proteomic features corroborate and extend previous observations, supporting the concept that a well-primed ASC secretome delivers a broad spectrum of bioactive molecules necessary for orchestrating effective peripheral nerve regeneration.

The regenerative capacity of peripheral nerves is sustained by the remarkable plasticity of SCs, which transition to a dedifferentiated repair state (repair SCs) following physical injury or neuropathic conditions ^34–37^. Upon nerve damage, SCs initiate regeneration by first dedifferentiating and proliferating in response to axonal signals, followed by redifferentiation and remyelination of regenerating axons ^38^. This reprogramming involves downregulation of myelin-associated genes and activation of reparative mechanisms, including secretion of trophic factors and cytokines, induction of autophagy, recruitment of macrophages for debris clearance, and formation of Bungner’s bands—specialized pathways that guide axon regrowth^39^. Unmyelinated axons, less than 1 μm in diameter, are grouped into Remak bundles, where each fiber is individually enveloped by non-myelinating SCs, or Remak cells ^40,41^. Both myelinating and non-myelinating SCs are crucial for maintaining the structural and functional integrity of peripheral nerves ^42^.

In the *in vivo* assessment, we have observed at different neurophysiological, molecular and morphological levels that in the groups treated with human and rat secretome this regenerative pattern has developed. A cornerstone finding of this research is the remarkable neurophysiological recovery observed in secretome-treated groups. Specifically, the evaluation of neurophysiological parameters, including the CMAP and NAP, demonstrated a significantly superior restoration of these potentials in the groups treated with secretome from both rat and human sources. This finding indicates a robust functional recovery and effective reinnervation following secretome application. Electrophysiological recordings, such as CMAP, are well-established methods for evaluating nerve regeneration after sciatic nerve transection injuries, alongside muscle weights and contractile forces ^43^.

These neurophysiological improvements are strongly correlated with the observed molecular and morphological changes. At the molecular level, the application of human secretome significantly increased NGF levels at the injury site. This finding aligns with the known neurotrophic properties of MSC secretome, which can increase levels of NGF and Brain-Derived Neurotrophic Factor (BDNF). NGF is a crucial neuroprotective and neurostimulatory factor that promotes regeneration following nerve trauma ^44^. qPCR data showed higher expression of myelin-associated genes (*MPZ, PERIAXIN*) post-injury, followed by their reactivation in secretome-treated groups, mimicking the SC dedifferentiation-redifferentiation cycle described by Jessen and Mirsky ^31^. MPZ and PERIAXIN are the most abundant myelin proteins in peripheral nerves, typically comprising 50-70% of total myelin proteins, and serve as a critical adhesion molecule for myelin sheath assembly and compaction. In severe nerve injuries such as neurotmesis, substantial myelin loss and *MPZ* and *PERIAXIN* downregulation would be anticipated as we could observe in the expression levels in neurotmesis group ^45^. However, confocal analysis revealed maintained MPZ protein levels across all experimental groups. This observation can be attributed to the remarkable stability of MPZ once incorporated into the compact myelin structure. The half-life of myelin proteins is considerably longer than their corresponding mRNAs, with MPZ protein persisting in myelin sheaths for weeks to months even after transcriptional arrest ^46^. Although the sustained elevation of pro-myelinating gene markers such as *Krox-20* and *c-Jun* is critical for effective myelination and efficient nerve impulse transmission, in our study, expression levels of these genes were highest in the biomaterial group, whereas the human secretome group exhibited values comparable to the control. This suggests that while secretome treatment supports a return to homeostatic gene expression associated with mature nerve function, the biomaterial alone may induce a more pronounced, possibly compensatory, upregulation of these markers in response to injury ^47,48^. *Krox-20*, for instance, is known to control myelination in the peripheral nervous system, and its upregulation is indicative of a healthy myelinating environment. In injured nerves, *c-Jun* also induces dedifferentiation of myelinating SCs to their immature form required for nerve regeneration ^49^. These molecular events collectively contribute to the observed structural and functional recovery ^50,51^.

The histomorphometry analyses in this study provide compelling evidence of near-complete nerve regeneration following secretome treatment, characterized by well-organized axons, robust myelin sheaths, and functionally intact Remak bundles containing properly ensheathed unmyelinated fibers (Figure 8D-E). These ultrastructural improvements align with the established role of SCs as key factors of peripheral nerve repair ^34,48^. The observed pro-myelinating SC activity and optimal vascularization reflect the secretome’s capacity to reactivate the SC repair phenotype—a dedifferentiated state essential for axonal guidance and myelin clearance. This finding aligns with ^37^Jessen and Mirsky’s work on SC plasticity ^39^, where successful regeneration requires SCs to transiently abandon their mature state to support repair processes.

The significant increase in myelin thickness within secretome-treated groups holds clinical relevance. Myelin thickness directly determines nerve conduction velocity, with thicker sheaths enabling faster signal propagation. Our measurements revealed a 2.3-fold increase in myelin thickness compared to untreated neurotmesis, explaining the restored CMAP amplitudes observed electrophysiologically. This structural-functional correlation mirrors findings by Chen et al. and Yu et al. ^52,53^, where MSC-derived exosomes enhanced remyelination in sciatic nerve injuries, correlating with improved conduction velocities.

Despite the promising results, certain limitations should be considered. While the rat sciatic nerve model is widely accepted due to its anatomical similarities to human nerves and its consistent regenerative capacity, translation to human clinical practice requires further investigation in larger animal models and eventually human trials. The small sample size, while justified by ethical considerations and the controlled nature of the experimental design, could benefit from expansion in future studies to further validate the statistical significance of observed differences and reduce potential biases. Future research should further investigate the optimization of secretome delivery methods to ensure prolonged therapeutic effects and characterize the precise molecular cargo responsible for specific regenerative processes. Elucidating the correlation between structural and functional recovery through multimodal approaches is essential for advancing new nerve therapies.

## 5. Conclusions

This study provides compelling evidence that ASC secretome, significantly enhances both structural and functional recovery following critical peripheral nerve injuries, as demonstrated by superior neurophysiological potentials, robust axonal regeneration, improved myelination, and favorable functional outcomes. The findings underscore the therapeutic potential of secretome as a cell-free biological strategy for nerve repair, offering a promising alternative to conventional methods and opening the way for future clinical translation.

## Data availability statement

Data will be made available on request.

## Declaration of generative ai in scientific writing

During the preparation of this work the authors used PERPLEXITY to improve readability and language of this manuscript. After using this tool, the authors reviewed and edited the content as needed and take full responsibility for the content of the publication.

